# Measuring motion-to-photon latency for sensorimotor experiments with virtual reality systems

**DOI:** 10.1101/2022.06.24.497509

**Authors:** Matthew Warburton, Mark Mon-Williams, Faisal Mushtaq, J. Ryan Morehead

**Author notes:** Correspondence concerning the article should be addressed to Matthew Warburton, University of Leeds, LS2 9JT. **Author Note**.

## Abstract

Consumer virtual reality (VR) systems are increasingly being deployed in research to study sensorimotor behaviours, but properties of such systems require verification before being used as scientific tools. The ‘motion-to-photon’ latency (the lag between a user making a movement and the movement being displayed within the display) is a particularly important metric as temporal delays can degrade sensorimotor performance. Extant approaches to quantifying this measure have involved the use of bespoke software and hardware and produce a single measure of latency and ignore the effect of the motion prediction algorithms used in modern VR systems. This reduces confidence in the generalisability of the results. We developed a novel, system-independent, high-speed camera-based latency measurement technique to co-register real and virtual controller movements, allowing assessment of how latencies change through a movement. We applied this technique to measure the motion-to-photon latency of controller movements in the HTC Vive, Oculus Rift, Oculus Rift S, and Valve Index, using the Unity game engine and SteamVR. For the start of a sudden movement, all measured headsets had mean latencies between 21-42ms. Once motion prediction could account for the inherent delays, the latency was functionally reduced to 2-13ms, and our technique revealed this reduction occurs within ∼25-58ms of movement onset. Our findings indicate that sudden accelerations (e.g. movement onset, impacts and direction changes) will increase latencies and lower spatial accuracy. Our technique allows researchers to measure these factors and determine the impact on their experimental design before collecting sensorimotor data from VR systems.

---

The recent development of low-cost consumer systems has seen virtual reality (VR) systems increasingly used to study behaviour. The potential for the use of VR systems in behavioural research has long been recognised. These systems offer the ability to present scenarios with high degrees of ecological validity (Loomis et al., 1999) while also allowing a level of control that goes beyond what is possible in the real world (Wann & Mon-Williams, 1996). This means users can be immersed in highly realistic simulations, but be asked to catch a ball that does not obey the laws of gravity (Fink et al., 2009) or explore non-Euclidean environments (Warren et al., 2017). This versatility offers a powerful tool for interrogating behaviour – from studying rodents, flies, and zebrafish (Holscher, 2005; Stowers et al., 2017) to examining the complexities of human psychological processing. Research into sensorimotor behaviour is increasingly harnessing these benefits, allowing complex sensorimotor skills such as golf putting and pool shooting to be studied with ease (Haar et al., 2020; Harris et al., 2020).

Popular use of the term VR now almost exclusively refers to head mounted displays (HMDs), a display mounted close to the eyes presenting stereoscopic images to give a sense of depth and allowing interactions via hand-held controllers. While historically expensive (Brooks, 1999; Slater, 2018), HMDs such as the Oculus Rift (Facebook Technologies, 2021) and HTC Vive (HTC Corporation, 2021) have made using VR in research a realistic option for many laboratories, not just those with specialist facilities. These HMDs have strong integration with popular game engines such as Unity (Unity Technologies, 2021) and Unreal Engine (Epic Games, 2021), which are used to develop professional VR games, and tools such as SteamVR (Valve Corporation, 2021) allow developed experiments to be deployed with any HMD. With the increasing use of these HMDs in behavioural research, a range of software frameworks have been created that specifically integrate with Unity to allow researchers to design experiments more easily and incorporate features common to behavioural research (Bebko & Troje, 2020; Brookes et al., 2019; Watson et al., 2019).

While consumer HMDs have been adopted to study behaviour, they were not designed to be scientific tools. As such, certain properties of these systems need to be verified to ensure the measurements they provide adequately address the research questions without confounding biases. One area where verification has thus far been lacking is in measuring the latency between the execution and visual feedback of movements, known as motion-to-photon latency (Figure 1a). The VR environment has many processing steps, each of which introduce latencies. These steps, shown in Figure 1b, include sampling the sensors, transferring the data to the computer and processing it, using the data to simulate a virtual environment, rendering the virtual environment to images, and displaying them on the HMD. The sum of delays along this pipeline gives the motion-to-photon latency. Because of these delays, by the time current tracking data is displayed on the HMD it is likely the HMD and controllers will be in different positions and orientations. Latency is known to affect end-user experience in VR, increasing levels of cybersickness (DiZio & Lackner, 2000) and lowering user’s feeling of “presence” in the environment (Welch et al., 1996). Much effort has therefore been dedicated to reducing latency in consumer VR systems (Carmack, 2013). Motion prediction is commonly used in consumer VR (LaValle et al., 2014) to reduce the effective latency of the system. This works by predicting the trajectory at the time the rendered images are presented to the HMD, rather than using the tracking data captured when the images are created. The effect of motion prediction is illustrated in Figure 1c, where the start of the movement is initially delayed. However, as soon as the motion prediction algorithm detects motion, it can extrapolate this to predict where the HMD or controller will be at the time feedback is presented, functionally reducing latency.

**Figure 1.**
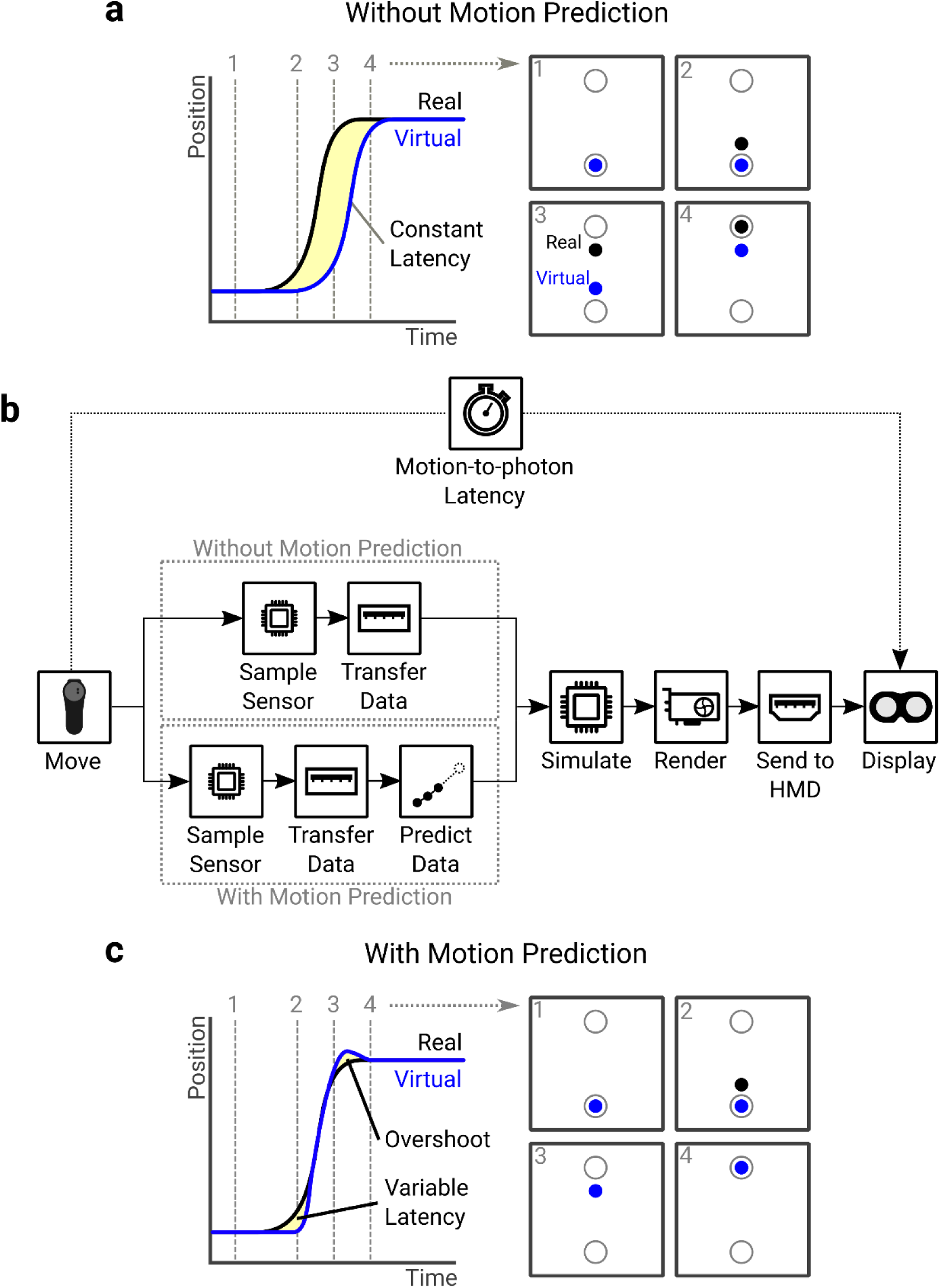
**(a)** Latency represented as a temporal difference between a real movement and its virtual visualisation. The effect of latency is shown in a reaching task, where the virtual movement lags behind the real one. **(b)** The pipeline of operations a position sample goes through to be presented to the user. Each step along the pipeline has its own latency, and the sum of these operations gives the motion-to-photon latency of the system. Motion prediction can be used to predict where the controller will be when the frame is presented to the user, functionally reducing latency. **(c)** Once a movement can be predicted reliably the virtual movement can match the real one. The effect of this is shown in a reaching task, where the initial portion of the movement is still delayed but once motion can be predicted the virtual movement matches the real one. The end of the movement may be similarly affected, with the virtual motion overshooting motion offset due to the deceleration.

It is important to understand the latency of experimental equipment because it has been observed to impact users’ sensorimotor performance. For instance, simple self-paced tasks such as handwriting and tracing are degraded in the presence of increased latency (Kalmus et al., 1960; Smith et al., 1960), with delays as little as 40ms increasing errors and time taken and reducing neatness (Bergeijk & David, 1959). Similarly, performance on manual tracking tasks worsens when an additional delay is introduced (Foulkes & Miall, 2000; Langenberg et al., 1998; Miall et al., 1985; Miall & Jackson, 2006; Smith, 1972; Vercher & Gauthier, 1992) with delays as small as 17ms enough to degrade the amount of time participants can lock on the moving target (Smith, 1972). While some studies demonstrate participants could adapt to these introduced delays (Foulkes & Miall, 2000; Miall & Jackson, 2006), temporal lags have been shown to disrupt the ability to adapt to visual displacements of hand movements (Brudner et al., 2016; Held et al., 1966; Held & Durlach, 1989; Honda et al., 2012; Kitazawa & Yin, 2002; Tanaka et al., 2011), with adaptation to prism displacement being reduced by delays as little as 50ms (Kitazawa et al., 1995). It is therefore critical that researchers determine the latency of the system used in their experiments, and ensure their conclusions are not undermined by this potentially confounding factor.

Previous studies investigating latency in HMD VR systems have thus far focussed only on movements applied to the HMD itself. Rotations applied to the Oculus Rift DK2 show latencies in the range of 40-85ms (Chang et al., 2016; Feldstein & Ellis, 2020; Raaen & Kjellmo, 2015; Seo et al., 2017), though a minority of studies have found latencies in the region of 1-26ms depending on the settings (Kijima & Miyajima, 2016; Zhao et al., 2017). Translations applied to the HTC Vive and HTC Vive Pro have shown latencies of about 22ms (Jones et al., 2019; Niehorster et al., 2017; Xun et al., 2019). Estimated values for other popular headsets, like the Oculus Rift CV1, Oculus Rift S, and Valve Index, are currently absent from the literature. Estimations for controller movements are entirely absent.

Several techniques have been used to assess motion-to-photon latencies. Pendulums or servo motors can be employed to find the difference in time between some motion feature like reaching the neutral point (Liang et al., 1991; Mine, 1993), zero velocity (Friston & Steed, 2014), or passing a threshold angle (Papadakis et al., 2011) or position (He et al., 2000). Some techniques have employed sudden movements, and found the difference in time between the actual movement onset and an indication that virtual motion had started (Bryson & Fisher, 1990; Feldstein & Ellis, 2020; Raaen & Kjellmo, 2015; Seo et al., 2017; Yang et al., 2017). Others examined the whole movement profile by comparing the real movement and a virtual representation, calculating some measure of average latency (Adelstein et al., 1996; Becher et al., 2018; Di Luca, 2010; Gilson & Glennerster, 2012; Scarfe & Glennerster, 2019; Steed, 2008; Zhao et al., 2017).

A critical issue with this variety of techniques is that they will produce different latency measurements if motion prediction is used. For sudden movements, where recent tracking data shows no consistent trend, the improvements from motion prediction are nullified. For continuous movements, like those produced by a pendulum, tracking data will show a consistent trend and motion prediction will reduce the effective latency. In the presence of motion prediction used on current HMD VR systems, we should expect the latency of sudden movements to be higher than continuous movements. A complete understanding of the system’s latency properties can only be gained by considering how latency changes over the full motion profile, as this latency changes dynamically over the course of a movement. The measurements are also dependent on the pipeline used to produce the virtual environment, for example what software is used to simulate and render the environment. An approach is therefore needed that applies to typical research setups and that allows accurate like-for-like comparisons between different VR systems.

We developed a novel latency measurement technique, where a 240fps smartphone camera recorded the movement of a VR controller mounted in a linear guide assembly (restricting motion to one plane) and the screen of an HMD. Simultaneously, a tool developed in Unity controlled the colour of the HMD screen in a predictable pattern and measured the virtual controller position. Video processing produced a file of real controller positions and HMD screen colours, which was then matched with a file output from Unity relating the virtual controller position to the HMD screen colour. An automated frame counting procedure was used to find the difference in time between real motion events and those events being reported to the HMDs. This technique was used to measure the input latency of popular immersive VR systems (HTC Vive, Oculus Rift CV1, Oculus Rift S, and Valve Index). To ensure the latency measurements are relevant to typical research setups, the Unity game engine and the Unity Experiment Framework (Brookes et al., 2019) were used to create the program that monitored the controller position and controlled the HMD screen’s colour and SteamVR (Valve Corporation, 2021) was used to ensure the input was HMD-agnostic. To account for the use of motion prediction we measured latency at two points – at the start of the movement, where motion prediction is nullified, and at the middle of the movement, where motion prediction can functionally reduce latency. Further, this technique allowed us to observe how quickly this reduction happens by assessing the latency across the whole movement.

## Method

### Latency Measurement Technique

A novel latency measurement technique was developed to assess the latency of the VR controllers, shown in Figure 2. A custom Unity program was created to monitor the position of a VR controller and control the screen of the VR HMD. The VR controller was mounted in a bench clamp which was connected to a linear guide, allowing the position of the VR controller to be varied in a single axis, and a battery powered LED was secured to this assembly. The colour of the HMD’s screen was controlled throughout, and a file was output for each trial with the virtual controller position and HMD screen colour at each time-step, sampled at the HMD’s frame rate. A smartphone was used to record a separate video of each trial at 240fps, ensuring the LED and the HMD were visible throughout (Figure 2a). A trial, shown in Figure 2b, was started by moving the controller assembly to either end of the linear rails and pressing a button on the controller, which turned the screen cyan for one HMD frame, blank for 50 frames, and then cyan for one more frame on each button press. After this the screen alternated between cyan and red every frame, apart from every tenth frame where the screen turned magenta.

**Figure 2:**
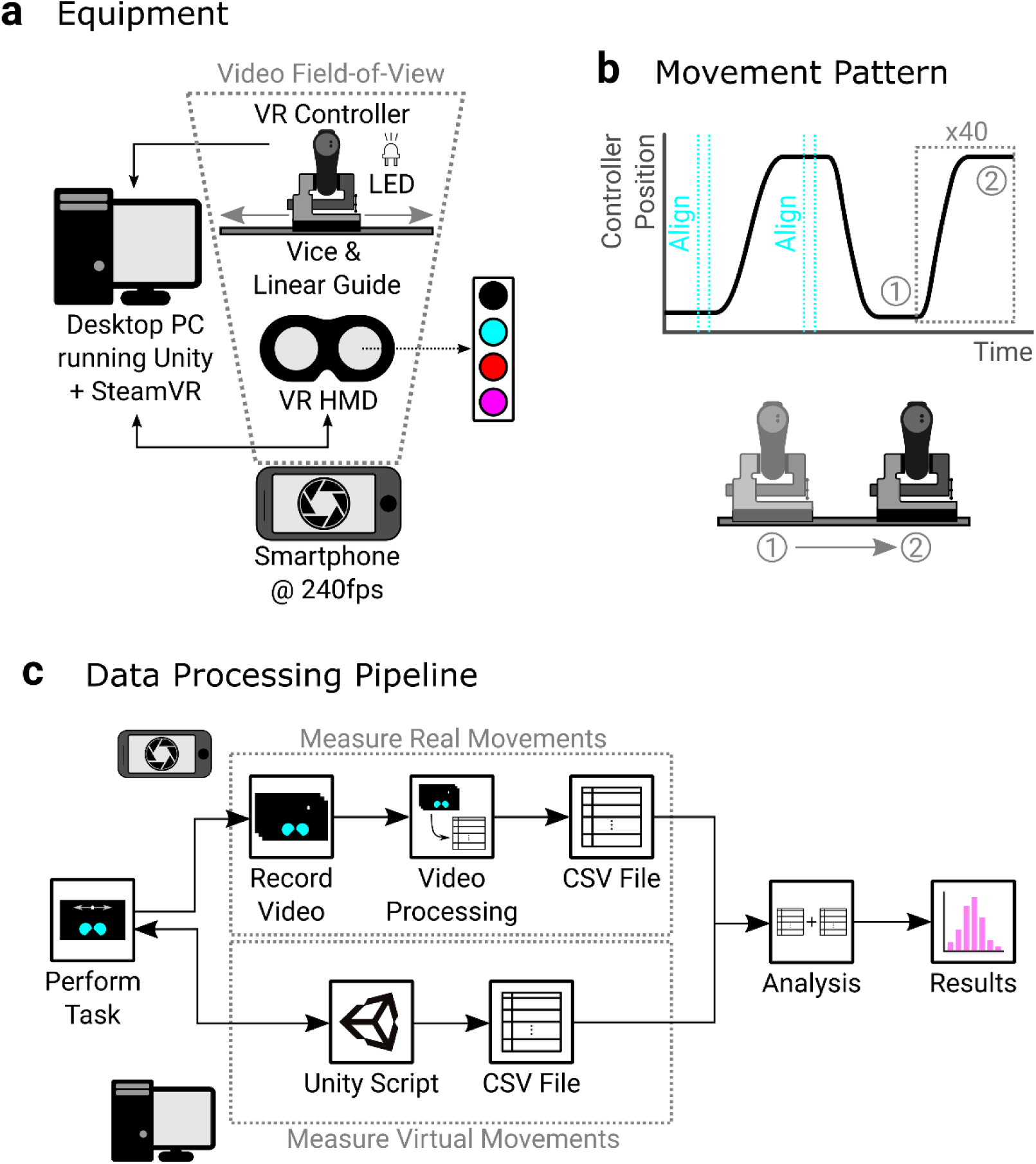
**(a) Equipment:** The experimental setup used to measure the input latency of VR controllers. A desktop PC ran a custom program that monitored the controller position and controlled the VR HMDs screen colour. The VR controller had an LED attached that was always on. A smartphone recorded the VR HMD and controller setups so that the LED and HMD were visible. **(b) Movement Pattern:** The controller assembly was moved to the extreme positions of linear rails and a button pressed to align the extreme positions, turning the screen cyan for one frame at the start and end of each alignment. Forty sudden movements were then applied to the controller assembly, translating it from one side of the linear rails to the other, with periods between movements to allow the equipment to settle. The screen colour after alignment was alternated between cyan and red every frame, apart from every tenth frame where the screen turned magenta. **(c) Data Processing Pipeline:** The Unity script controlled the HMD screen colour through the task and recorded the VR controller position and screen colour at every HMD frame, and output this to a CSV file. While the task was being performed, a video was recorded of the HMD screen colours and real controller position. Video processing was performed on the videos to convert them to a CSV file containing the HMD screen colour and real controller position at each camera frame. An analysis script used both output files and matched camera and HMD frames where the screen colour was shared, allowing the real and virtual controller positions to be linked. The analysis script then used these matched real and virtual controller positions to measure the motion-to-photon latency at different parts of the movement.

Pilot testing of detecting the HMD screen colour showed that this combination of colours allowed the highest discriminability once recorded on the smartphone, such that frame colour was never mis-classified. A series of sudden movements followed by a settling period were then applied to the VR controller assembly, moving it from one end of the linear rails to the other. Some techniques suddenly shove the device being measured with a hand (Niehorster et al., 2017). We wanted to ensure the movement onset was sudden enough to detect accurately, so our sudden movements were initiated by hitting the side of the VR controller assembly, after a short backswing, with a blunt object (the handle side of a screwdriver in this case) and then pushing the assembly to the opposite end of the rails. Forty such movements were applied to the controller assembly per trial by the experimenter. To standardise the movements between systems, the movements were applied at a rate of approximately 1 movement cycle per 3 seconds (around 0.75s to move, and 2.25s to settle).

Our technique used the HMD screen colour as an intermediary to link the real and virtual controller positions. By matching the screen colours in the file output from Unity to the frames in the video, the real and virtual controller positions could be co-registered. The sequence of operations is shown in Figure 2c. The videos were processed offline to give files containing the real controller position and the colour of the HMD screens at each camera frame. The analysis took the file output from both the video processing and Unity for each trial and matched the camera frame where the HMD first changed to a new colour to that observation in the Unity file. As the camera sampling rate was higher than that of the HMD, one HMD frame lasted multiple camera frames, so we were only interested in the first camera frame where a certain colour was shown. This link between the real and virtual screen colours allowed the virtual controller position to be matched to real controller positions, as detailed in the Analysis section. Latency was then assessed at the start and middle of each movement, to mimic the typically used techniques, and across the full motion profile, to see how quickly motion prediction allows the latency to be functionally reduced.

### Equipment

#### Desktop PC

The program used to interface with the VR systems was run on a desktop PC with the Windows 10 operating system. The computer specifications were an AMD Ryzen 2600X CPU, 32GB DDR4 RAM, and Nvidia GTX 1060 GPU (driver version 457.09), meeting at least the minimum specification for all HMDs tested.

#### VR System Assembly

The latency of four VR systems was assessed: HTC Vive (1080 x 1200px per eye, 90Hz), Oculus Rift CV1 (1080 x 1200px per eye, 90Hz refresh rate), Oculus Rift S (1280 x 1440px per eye, 80Hz), and Valve Index (1440 × 1600px per eye, 80Hz/90Hz/120Hz/144Hz). As the Valve Index features four different frame rates, in total seven VR setups were tested. These headsets were chosen as they are flagship systems from the most popular manufacturers of VR HMDs and dominate the share of devices used for gaming (Lang, 2020). The HMD and right-hand controller for each system were used when assessing latency. All the tested HMDs feature low pixel persistence where the screen is only illuminated for a short period at the end of each frame to reduce motion blur (0.33ms for the Valve Index – 2ms for the Oculus Rift). For the Oculus Rift this means that for a 11.1ms frame duration, the screen is blank for 9.1ms, and only illuminated for the final 2ms. The captured videos therefore have frames where the HMD screen is not illuminated between frames where it is illuminated (as an example, Figure S1 in Supplementary Materials shows the screen turning black, instead of another colour, when the HMD frame swaps).

A bench clamp (Panavise 301, 1.2kg) was used to hold the VR controller and was connected to a linear rail assembly allowing 185mm of travel which compromised two Igus TS-04-15-300 linear guide rails and four Igus TW-04-15 T linear guide blocks. This assembly allowed a common setup to be used for all VR systems and restricted movement of the controller to a single axis which was orthogonal to the view of the camera, maximizing the camera’s sensitivity to measure controller position. An LED powered by a CR2032 battery was secured to the vice, close to the controller.

#### Smartphone

A Google Pixel 4a smartphone (Android 11), mounted on a tripod, was used to record the videos. The videos were recorded at a resolution of 1280x720 pixels and a framerate of 240fps. The default camera settings for the phone were used when recording the videos, beside brightness being lowered to improve contrast. Recordings were started remotely using volume control buttons on a pair of earphones plugged into the phone via the 3.5mm audio jack. To validate the framerate of the camera, a microcontroller (Elegoo Uno R3) was used to turn on an LED every second for 30 seconds, and a video was recorded to count how many frames passed between two subsequent flashes. This gave an average framerate of 239.90fps, negligibly different than the nominal framerate (0.002ms difference in frame length).

### Software

#### VR Program

A custom program was written to perform the VR elements of the experiment. To ensure the latency measurements would be representative of typical VR experiments, the Unity (version 2019.4 LTS) game engine (Unity Technologies, 2021) and SteamVR plugin (version 1.14.16, Unity plugin version 2.6.1) (Valve Corporation, 2021) were used to develop the program. The SteamVR plugin ensures that VR input to the program is agnostic of the specific HMD used. Default SteamVR settings were used for each HMD. The Unity Experiment Framework (Brookes et al., 2019) was used to ease program development and output a file for every trial containing the screen colour and virtual controller position at every HMD frame. Prior to testing each setup (including different frame rates for the Valve Index), the built-in software was used to calibrate the tracking and set the origin and forward-facing direction (Oculus SDK for Oculus Rift and Oculus Rift S, SteamVR for HTC Vive and Valve Index).

In the program, the 3D position of the VR controller was monitored just before the frame is rendered, where SteamVR updates the controller position. The colour of the HMD screens was controlled by covering each screen’s rendering camera with textures representing the different available screen colours (black, cyan, red, magenta) and using culling masks to only display the required colour on a given frame. The screen colour update was triggered by SteamVR updating the controller position. A file was output every trial giving the position of the controller and the colour of the HMD screens at every HMD frame. As Unity recommends no additional computation be done in the loop once the controller position is updated, these data were gathered in the late update loop. This loop runs before the controller and screen are updated, meaning the reported position and screen colours lag the time stamp by one frame, which was corrected for in the analysis.

#### Video Processing

A semi-automated video processing procedure was performed using Python (version 3.7.0) and the OpenCV (version 4.2.0) image processing library (Bradski, 2000). For a folder of videos, this procedure searched for the position of the LED and colour of the HMD screen within user-identified regions of interest (ROI) around the controller assembly LED and within the HMD screens. This allowed the real and virtual movements to be co-registered for each video frame. The high-level description of this data processing pipeline is shown in Figure 3a.

**Figure 3:**
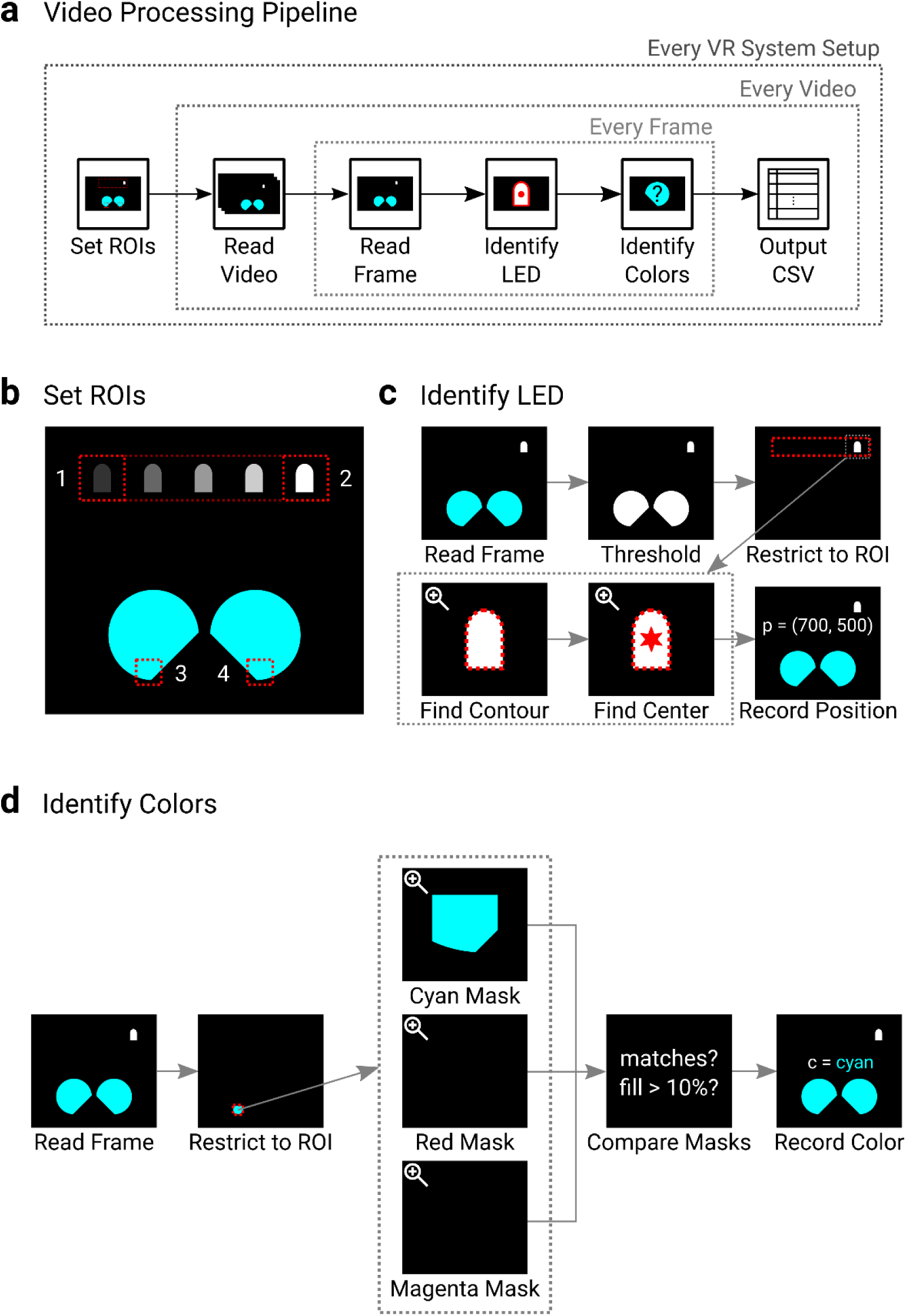
**(a) Video Processing Pipeline:** For each VR system setup, the first video is used to identify the regions of interest (ROIs) in the video frames. After this, each frame of each video for that setup is probed to find the LED and screen colours. The results for each video are output to file. **(b) Set ROIs:** On the first video for each setup, four positions are manually identified by the user: (1) controller left extreme position, (2) controller right extreme position, (3) left screen, and (4) right screen. **(c) Identify LED:** On each frame the LED is found by converting the image to black-and-white via a threshold, restricting the video to only pixels bounded by the rectangle covering ROIs 1 and 2, finding the largest contour in the frame and then identifying the centre of the contour, which is recorded as the controller’s real position. **(d) Identify Colours:** On each frame the screen colour is found by restricting the video frame to only pixels inside either ROI 3 or 4 (to find left or right screen colour respectively) and creating three separate masks indicating whether the pixels are either cyan, red, or magenta. Checks are then performed on the masks. If no masks have any matches the screen colour is recorded as black. If a mask does have a match and more than 10% of the ROI is filled then the screen colour is recorded as the mask with the most matches, and if not is recorded as being ambiguous, and can be corrected for in post-hoc analysis. This is performed for both the left and right screens.

During the processing of each folder of videos, user input was required to identify the extreme positions of the controller assembly LED and a point in each HMD screen. A window with a slider to control the frame number was presented, and left mouse clicks were required at roughly the centre of the LED in the two extreme positions. A left mouse click was required at a point in each HMD screen, at a point low enough in the screen that the ROI would include the bottom edge (an ROI was used to reduce video processing time and restrict analysis to only regions where the screen should be, and a point on the lower edge was required as the camera’s rolling shutter meant pixels at the bottom would be the first to illuminate). A second window with a slider to control the threshold (on a scale of 0-255) used when converting frames to black-and-white was then presented. A threshold value of 150 was used throughout as pilot analysis indicated this adequately captured the LED outline. This process was performed only on the first video in the supplied folder of videos. The user-supplied positions were used to define an ROI around the possible VR controller assembly LED positions by taking the line between the left and right alignment positions and padding in all directions by 20 pixels to account for the height and width of the LED, giving a rectangle where the LED should always be found. An ROI was then defined for each HMD screen by padding the selected points by 20 in all directions. The same ROIs were used for every video inside the folder. The process of identifying the ROIs is shown in Figure 3b.

The rest of the process was then automated for every video inside the folder, performing the same steps on each frame of each video. To find the position of the VR controller assembly LED on each frame, the image was turned to black and white using the user-supplied threshold, and the LED ROI was then used to turn any pixels outside of it black, so only white pixels related to the LED should remain.

Any contours of white pixels were then identified, and if multiple existed the largest was taken to be the LED. The position of the centre of the contour (rounded to the nearest pixel in both dimensions) was then identified and stored. The steps to find the LED are shown in Figure 3c.

To find the colour of each HMD screen, the pixels inside each screen ROI were first converted to hue saturation value (HSV) coordinates, and three separate image masks (one for each colour) were created to indicate whether the pixels in the ROI matched the following criteria:

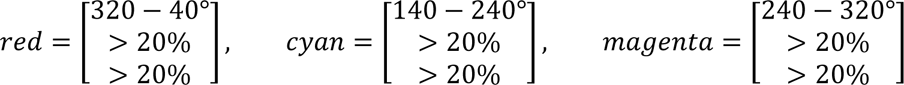

As black screens had very low saturation and values (typically zero for each), the checks on these properties ensured only coloured pixels were matched. Because only three colours were used, they were each given a wide hue range to ensure there was little chance of coloured screens not being detected. If no pixels were matched in any of the three colour masks, then the screen was classified as being black. If at least one mask was matched, then a check was performed on the mask with the most matched pixels to see if more than 10% of the ROI had matched pixels. If this was true, the screen colour was classified as the mask’s colour. If not, then the screen’s colour was marked as ambiguous, and it was checked whether the matched pixels were concentrated in the top or bottom of the ROI. The ambiguous classification was needed because the camera and HMD refresh were not synchronized, so some frames captured the HMD screen just turning on (where, due to the camera’s rolling shutter, the matches were concentrated in the bottom of the ROI) or off (where the matches were concentrated in the top of the ROI), leading to a low amount of the ROI pixels matching. Pilot testing showed colours were harder to distinguish when the screen was turning on or off. However, for such frames where a low amount of the ROI was filled due to the screen turning on or off, the camera’s rolling shutter guaranteed that the following or preceding frame respectively featured enough coloured pixels in the ROI to be accurate.

Therefore, marking the frame colour as ambiguous and indicating where pixels were concentrated allowed the screen colour to be corrected during post-hoc analysis. The steps to find the screen colours are shown in Figure 3d, and the handling of ambiguous frames is demonstrated in supplementary material (Figure S1).

These steps were then repeated for every frame. A comma-separated variable (CSV) results file was exported for each video with a row for each frame containing the frame number, the 2D coordinates of the VR controller assembly LED in pixels, and the colour of each HMD screen. A video was also exported that added this information to the original video so the results could be validated by adding a line representing the outline of the LED, a point representing the LED’s centre, and text above each screen indicating the detected screen colours.

To test the performance of the video processing procedure at reporting the correct colour, a script was created that used the first video output by the video processing script of each setup (with the identified screen colours added to each frame) and selected 500 random frames from each video to display to the validator. For each randomly selected frame, the validator was presented with the previous, current, and next frame displayed side by side, and had to indicate whether the screen colours of the current frame were either stated correctly (text agreed with screen colour), or where the screen colour was ambiguous due to partial illumination whether it could be identified by looking at the previous or next frame if the partial illumination was at the top or bottom of the ROI respectively. If the information was correct the validator pressed the ‘Y’ key, and otherwise pressed the ‘N’ key. This process was repeated until the randomly selected frames for each setup had been classified. This showed that all screen colours presented to the validator were either correct or could be identified during analysis.

## Analysis

The analysis script was written using R (version 3.5.3; R Core Team, 2020). The analysis script read in the CSV files created during the video processing, giving the real controller position and HMD screen colour for each camera frame, and the VR program, giving the virtual controller position and HMD screen colour for each HMD frame.

The first process performed was to co-register the real and VR movements by aligning the camera and HMD frames featuring the same screen colour. A reduced data set for the real controller movements was first created so that the virtual controller positions could be paired, which was then integrated back into the full data set. In total 2,107,500 camera frames (observations) were processed during the analysis. For each real result file, any missing positional data for the LED was spline interpolated (1 observation Valve Index at 120Hz, 182 observation Oculus Rift S). The HMD screen colour was corrected for any observations where the colour was marked as ambiguous, by either using the screen colour from the previous or next observation where the ambiguous colour was at the top or bottom of the ROI respectively. The data set at this point was the ‘full’ data set of real controller movements. As all the HMDs used low pixel persistence the camera captured black frames between the HMD screen being illuminated, so these were filtered out. A single screen colour was created for each observation by checking whether either screen reported a non-black colour. There were no cases where the HMD screen colours did not match each other. Because the HMD could be illuminated for multiple camera frames, only the first observation was kept where the same colour was reported for more than one frame consecutively, as we are only interested in the earliest time a particular frame colour was reported. For both files this left only observations where either the alignment procedure or main task was performed. The real and virtual alignment positions, to be used later in the analysis, were found by calculating the mean controller position during the two alignment procedures and these observations were then excluded to leave only observations from the main task. This gave the reduced data set used to co-register real and virtual movements.

With the pre-processing performed, the screen colours from the video processing and VR program then needed to be matched. Counters were initialised at the first observation for the real and VR movements and were iterated together until either the end of the real movement data was reached or there was a discrepancy between the real and VR screen colours. Discrepancies between the real and VR screen colours were rare (2 corrections Valve Index at 80Hz, 4 correction Valve Index at 120Hz, 4 corrections Valve Index at 144Hz, 12 corrections Oculus Rift, 2 corrections Oculus Rift S). Though impossible to confirm, these discrepancies resembled the camera either dropping or repeating a frame. Whenever a discrepancy was encountered, a process was performed to realign the real and virtual datasets at the next matching purple frame. If the real dataset reached a purple frame sooner than expected (likely camera dropped a frame) then the virtual counter was moved to the next purple frame, and the counters were iterated again. Otherwise, this was done by moving both counters back to the previous congruent purple frame, and then finding the difference in time between this frame and the next 50 purple frames for both data sets. A search was then performed to find the pair of real and virtual purple frames that had the closest time difference from the previous congruent purple frame. The counters were then set at these frames and iterated again.

In total these discrepancies lost 4 sudden latency measurements (2 for the Valve Index at 80Hz, 1 for the Valve Index at 144Hz, and 1 for the Oculus Rift) and 5 continuous latency measurements (2 for the Valve Index at 80Hz, 1 for the Valve Index at 144Hz, and 2 for the Oculus Rift). Further, one video of the Valve Index at 120Hz only featured 30 movements, reducing the number of sudden and continuous measurements for this video by 10. This process gave a link between the real and virtual frame numbers, which were then used to incorporate the virtual controller positions into the ‘full’ data set containing the real controller positions. This led to a data set that had real controller positions at every sample but only VR controller positions on the first frame of every screen colour alternation.

### Latency for sudden movements

To find the latency of the sudden movements, each movement onset needed to be identified. While it is typical to use methods like determining when a movement passes a threshold speed or percentage of peak speed to find reaction times (Brenner & Smeets, 2019), to ensure accurate measures of latency we need to know the first frame that movement started which requires a more precise method. A simple movement onset technique could be used to find the real motion onsets, as the resolution of the videos was relatively low (∼2.12px/mm), the real controller positions were affected little by noise and showed monotonic changes in position during movements. To determine movement onset, the mid-point of each end-to-end movement was found and at each of these frames the direction of movement (1 for increasing position, -1 for decreasing position) was defined. A backward search was then initiated at each mid-point, terminating when either the position difference to the previous frame was in the opposite direction to that expected (i.e. [𝑝_𝑓_ − 𝑝_𝑓−1_] × 𝑑𝑖𝑟𝑒𝑐𝑡𝑖𝑜𝑛 < 0), or in the case where the position difference was zero, where the position difference to the frame before that was zero or in the opposite direction (i.e. [𝑝_𝑓_ − 𝑝_𝑓−2_] × 𝑑𝑖𝑟𝑒𝑐𝑡𝑖𝑜𝑛 ≤ 0). This was then the first frame where motion thereafter was consistently in the same direction, while not misclassifying the controller as being stationary during pairs of frames where the controller position did not change (e.g. due to slow movements). As the initial motion was always sudden any pairs of frames with zero position difference should be further into the movement and should not affect motion onset detection.

To ensure the method captured real motion onset well, a sample of 40 randomly selected movement starts was manually classified for each setup. An application programmed in R presented the validator with a graph of the cursor position over time, restricted to the specific motion start, and the validator selected the data point where motion onset occurred. This was performed for each randomly selected movement for each setup, and the validator-selected points were compared to those automatically detected. The only discrepancies were for the Valve Index at 120Hz, where one motion onset was automatically identified 3 frames later than the manual classification, and the HTC Vive, where one motion was detected 6 frames later than manual classification. Assuming this rate of discrepancy was representative of the whole sample for that setup, it would have affected mean measured latency by about 0.3ms and 0.6ms respectively.

As the spatial resolution of the VR movements was higher than the video, noise meant that a stationary controller led to non-stationary position samples, so the same technique could not be used here. Instead, motion onset for the VR movement was classified using an outlier detection technique that compared each new position sample to the median absolute deviation (MAD) of a window of previous samples (Leys et al., 2013):

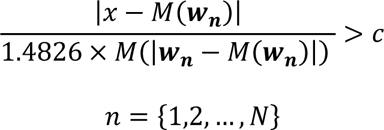

where 𝑥 is the new position sample, 𝑀 is the median function, 𝒘 is the window of previous positions, 𝑛 is the element number in window with length 𝑁, and 𝑐 is a threshold. The timestamp of the measured real controller onset, identified above, was used as a start point, and the frames featuring a VR movement were then iterated through forward until the inequality was met. This then was the first frame where movement deviated meaningfully from the previous samples, indicating movement had started.

Finding the optimal parameters for the outlier detection method involved two steps. The virtual motion start procedure was first run using a default pair of parameter values (𝑤 = 38, 𝑐 = 12.5) that provided reasonable detection during pilot testing. The same application used to validate real movement onset was used to present the validator with 40 of these virtual movement onsets for each setup. The validator then manually identified the motion onset. A grid-search was then performed where every combination of parameters in the range 𝑁 = [10, 11, …, 40] and 𝑐 = [5, 5.1, …, 20] were used to generate frames where motion onset would be detected, and these were compared to the manually classified onsets. Parameter values could be found that gave no misclassifications in the sample for the HTC Vive or the Valve Index (at all frame rates). The best performing values for the Oculus Rift had three misclassifications, where two observations were detected two frames early and one observation was detected two frames late, and for the Oculus Rift S had two misclassifications, where two observations were detected three frames earlier. Assuming this is representative of the whole sample, mean measured latency would be 0.2ms and 0.6ms lower than it should be for the headsets respectively. Virtual motion onset detection was then re-run with the identified optimal parameters for each setup. This validation procedure adds confidence that the motion onset algorithms work well, but as described, a small number of the samples were not perfectly detected. This is likely to lead to some individual samples providing lower or higher values than the true latency, but should have minimal impact on the group averages.

The real and VR movement onset frames were then matched together. A fuzzy left join was performed between the real and VR movement onset for both screens, where observations were matched if the absolute frame difference was up to 25 frames, and the difference in frames between the real and VR movement onset frames gave the latency. In the case where multiple matches existed for a detected movement onset the only the smallest difference was kept. The distribution of these differences was then visualised in histogram plots. This process was performed for each result, and the results for each trial were then collated and plotted together to give the overall latency distribution of the HMD. Values outside of 3 standard deviations of the mean value for each setup were discarded, removing 19 observations (2 observations Valve Index at 80Hz, 2 observations Valve Index at 90Hz, 5 observations Valve Index at 120Hz, 8 observations Valve Index at 144Hz, 2 observations HTC Vive). As both the screen illuminating and the real controller starting to move have uncertainty arising from the sample-and-hold nature of video frames (e.g. the real controller could have actually started moving at any point in the time between the current and previous frame), this will on average cancel out but give a ±4.2ms uncertainty around any single value (Feldstein & Ellis, 2020).

### Latency for continuous movements

To find the latency of continuous movements, the frames where the controller crossed threshold positions needed to be identified. The threshold positions were defined in relation to the alignment positions, so the real and VR positions were normalised by making zero and one represent the two alignment positions. The frames where the real and VR position crossed the mid-point between the two alignment positions were then identified (i.e., the mid-point lay between the current and previous position). The process to match real and VR mid-point crossing was the same as for the sudden movements. Values outside of 3 standard deviations of the mean value for each setup were discarded, removing 2 observations (1 observation Valve Index at 144Hz, 1 observation HTC Vive).

### Latency over the full movement trajectory

To understand how the latency developed after the sudden start, the complete real and virtual motions were compared. The first comparison was to find the latency at which each real controller position could be displayed on the screen of the HMD. To do this, all movements were aligned at motion onset and the positions were normalised about the two threshold positions as in the continuous movements, but also modified so all movements acted in a positive direction (i.e. any movements where controller position decreased were reversed). At each time point from the motion onset the normalised real controller position was compared to the normalised virtual controller positions to find the next time the HMD screen was on that could have reported this movement. This was done by projecting the normalised real controller position forward in time until it intersected a line between two normalised virtual controller positions, and then finding the next frame that the HMD screen turned on. The difference between these two frames was then classed as the minimum possible latency of that controller position given the constraints of the HMD frame rate. This was repeated for each normalised real controller position. This gives a measure of the latency for each real position sample, but because only the time that the HMD screen turns on has uncertainty due to the sample-and-hold behaviour, the measure will on average over-report latency by half a camera frame’s duration (i.e. 2.1ms). To also understand how this latency impacts the presentation of the controller position on screen, the difference in normalised position between the real and virtual controller positions was also calculated for each time step. This then gave the positional error that was induced by the latency of the system. Both measures allow the time course of the motion prediction to be assessed, as the latency and positional error should drop from movement onset, so the time when latency and positional error stabilise should indicate how long the motion prediction takes to “warm up”.

## Results

The novel latency measurement technique was used to assess the properties of the controller position input of four HMDs (Oculus Rift, Oculus Rift S, HTC Vive, Valve Index) in two movement conditions – Sudden Movement, where movement onset is detected, and Continuous Movement, where the controller was moved smoothly after the sudden motion onset, shown in Figure 4a.

**Figure 4:**
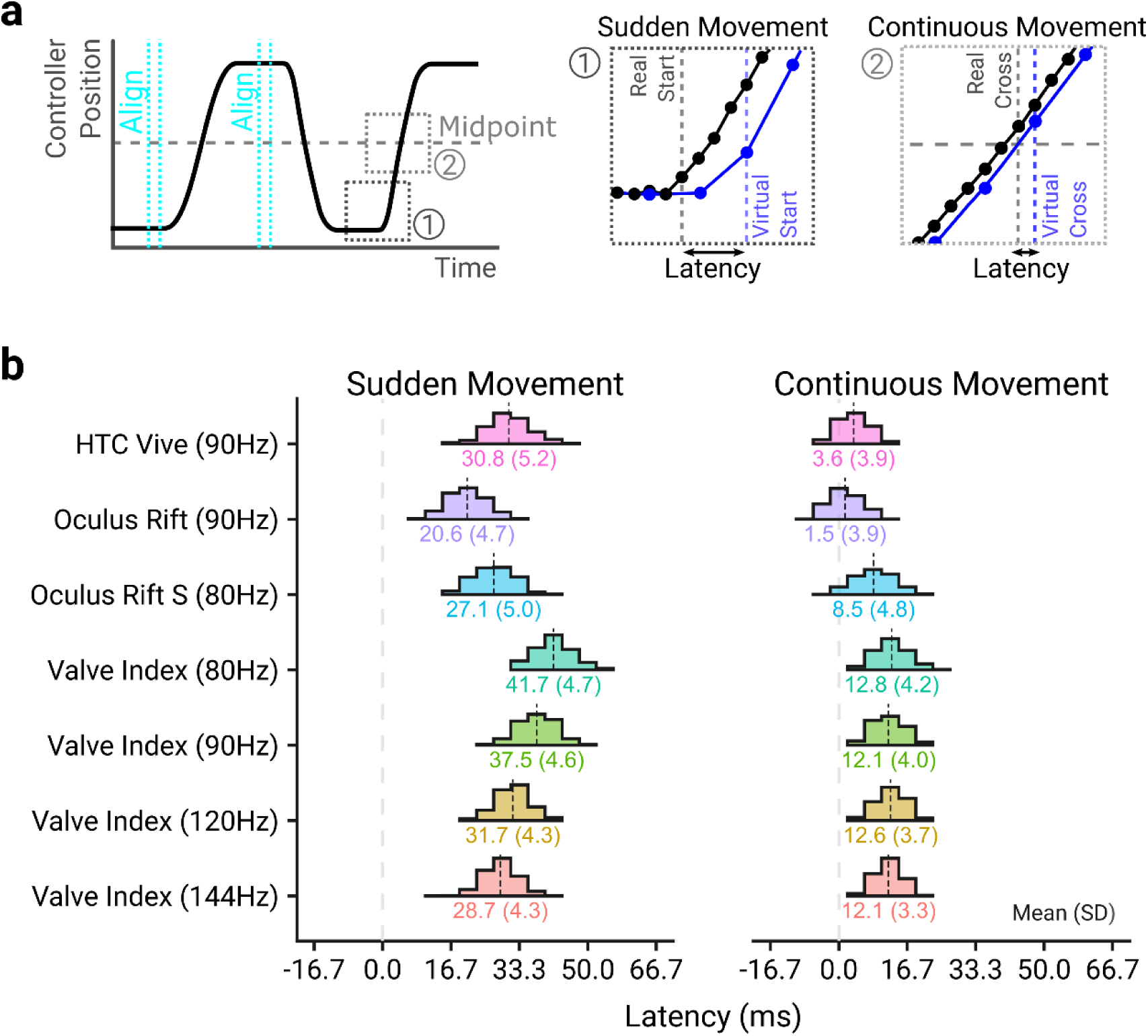
**(a)** The latency at the start (Sudden) and middle (Continuous) of the movement were measured. Note that the real controller positions were sampled every camera frame, whereas virtual controller positions could only be sampled every HMD frame. **(b)** Histograms of the measured latency for the different HMDs in the Sudden Movement (left panel) and Continuous Movement (right panel) conditions. The mean and standard deviation for each HMD and frame rate combination is shown beneath the histogram. The histogram bin widths were 4.17ms, to match the camera frame rate, centred on a latency of 0ms. Some of the variability present in the measurements is due to the stochasticity between the event occurring (the movement onset or the mid-point crossing), and when the camera captures a new frame or the HMD displays a new frame, as illustrated in Figure S2 in supplementary materials.

The time between a sudden movement being applied to the controller assembly and the movement being reported on the screen of the different HMDs is shown in the left panel of Figure 4b. This indicates that the HMD with the lowest latency was the Oculus Rift, and the HMD with the highest latency was the Valve Index operating at 80Hz. The Valve Index operating at 144Hz was only marginally better than the HTC Vive (90Hz) and worse than the Oculus Rift (90Hz) and Rift S (80Hz), despite having a much higher refresh rate than these HMDs. The measured latency was consistent between recordings for all HMDs, with the highest between-video standard deviation being 1.75ms for the Oculus Rift S.

While moving the controller suddenly ensures the measured latency is not affected by motion prediction, movements tend to be executed smoothly and as such, we expected lower latencies later in the movement, where accurate motion prediction was possible. The time between the movement mid-point being crossed by the controller assembly and the crossing being reported on the screen of the HMD is shown in the right panel of Figure 4b. It is clear from the comparison between Sudden and Continuous Movement conditions that motion prediction allows the base latency of the system to be ameliorated. Both the Oculus Rift and HTC Vive have near-zero latency once motion prediction is functioning, while the Valve Index, with the worst continuous latency, has mean latencies below 13ms across all refresh rates. Again, the latencies were consistent between videos, with the highest between-video standard deviation being 0.75ms for the Valve Index at 120Hz.

A linear model (R² = 0.89, F(13, 5,535) = 3,424.05, p < .001) was fit to the sudden and continuous latency data with effects of HMD setup and type of latency measurement. The model showed significant main effects of HMD setup (F(6, 5,535) = 1,202.99, p < .001), indicating the measured latency depended on the HMD setup, and measurement type (F(1, 5,535) = 35,813.28, p < .001), with sudden latencies being significantly higher than continuous latencies, as well as a significant interaction between HMD setup and latency type (F(1, 5,535) = 246.91, p < .001). Post-hoc comparisons using Bonferroni-Holm correction showed that all HMD setups showed significantly different latencies for sudden movements (p’s < .003), and that all HMD setups showed significantly different latencies for continuous movements (p’s < .001) except for comparisons between the different Valve Index frame rates which were all non-significant (p’s > .073). Further, all HMD setups showed a significant decrease in latency between sudden and continuous movements (p’s < .001).

As the sudden and continuous latency measurements only capture a snapshot of the latency properties, it is unclear how the latency develops over time. To understand this, the real and virtual motions were compared to understand how soon after motion onset the latency could be reduced by motion prediction. This comparison was performed for all HMD setups but only visualised for the Oculus Rift and the Valve Index at 90Hz, as these HMDs had the lowest and highest latency for that frame rate respectively. A comparison of the movements for the two HMDs is shown in Figure 5a. This shows there is an initial delay for the virtual controller to move, after which the virtual controller over-compensates to catch up to the real one before stabilizing. The systems also show overshooting behaviour upon a sudden stop, shown in Figure S3 in the supplementary materials, but the current experiment was not designed to assess this in detail. Note that Figure 5a shows, on average, that movements start earlier than the sudden latency measurements would suggest (e.g. the Valve Index position at 29ms is non-zero). This is due to finding the mean position of a small number of movements that have initiated and a larger number that have not, in accordance with the histograms reported in Figure 4b.

**Figure 5:**
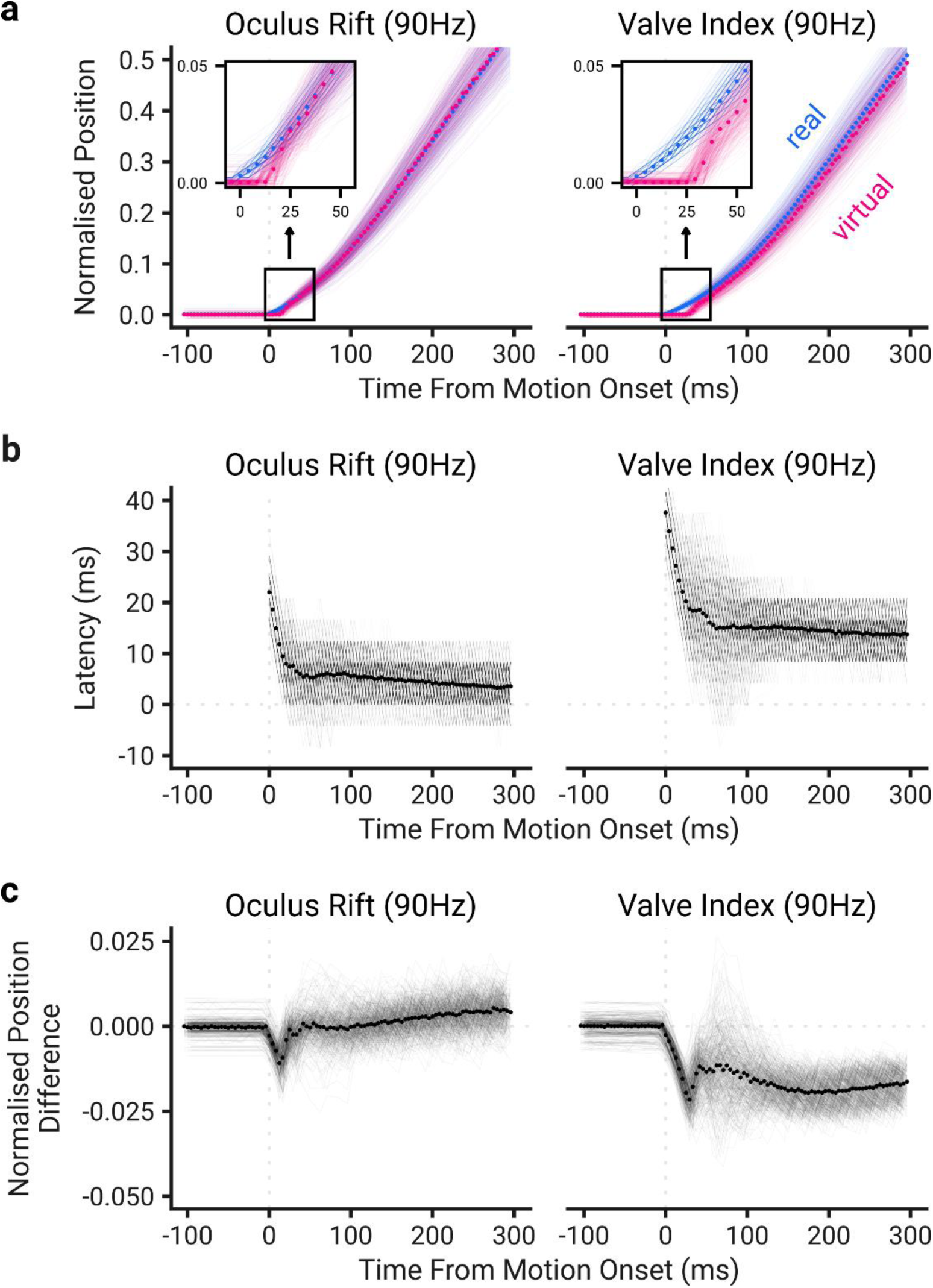
Latency properties of the Oculus Rift and Valve Index at 90Hz during a movement. Points show the mean value while the lines show individual movements. **(a)** Each movement was plotted relative to motion onset. Inset panels show a closer view of the motion onset. **(b)** The minimum latency that each real position could be displayed on the HMD was found. This was done for each sample after motion onset by projecting the real controller position forward in time until a greater virtual controller position was found, and then finding the next frame where the HMD was illuminated. **(c)** The difference between the virtual and real normalised positions was found at each time step to show the effect latency has on what is displayed by the HMD.

The real controller position at each time point was compared to the next time point that the HMD could reflect this position to give a measure of the latency over the entire movement (Figure 5b). This confirms the HMDs have high latency initially before reaching a plateau, which is reduced slowly through the rest of the movement. Note, however, that the shape of the latency curve will depend on the movement performed. We fit an exponential distribution to the latency values during the first 100ms of the movement to determine how quickly motion prediction would reach its ‘continuous’ latency. Specifically, this was where the difference in fitted values first fell below a standard deviation of the fitted residuals (where noise begins to outweigh signal), which indicated the latency stabilised at low levels within the first 33ms (Oculus Rift) to 54ms (HTC Vive) of the movement. We also tested a different method, when the latency falls within an arbitrary threshold of 2ms from the asymptotic value, which found similar values (25ms Oculus Rift to 58ms Valve Index at 80Hz). The latency over the first 100ms of motion is shown for all systems in Figure 6. As detailed in the methodology, this procedure will over-estimate latency by 2.1ms on average because the real controller position is used as a reference point, meaning the measurement uncertainty only affects the virtual controller position.

**Figure 6:**
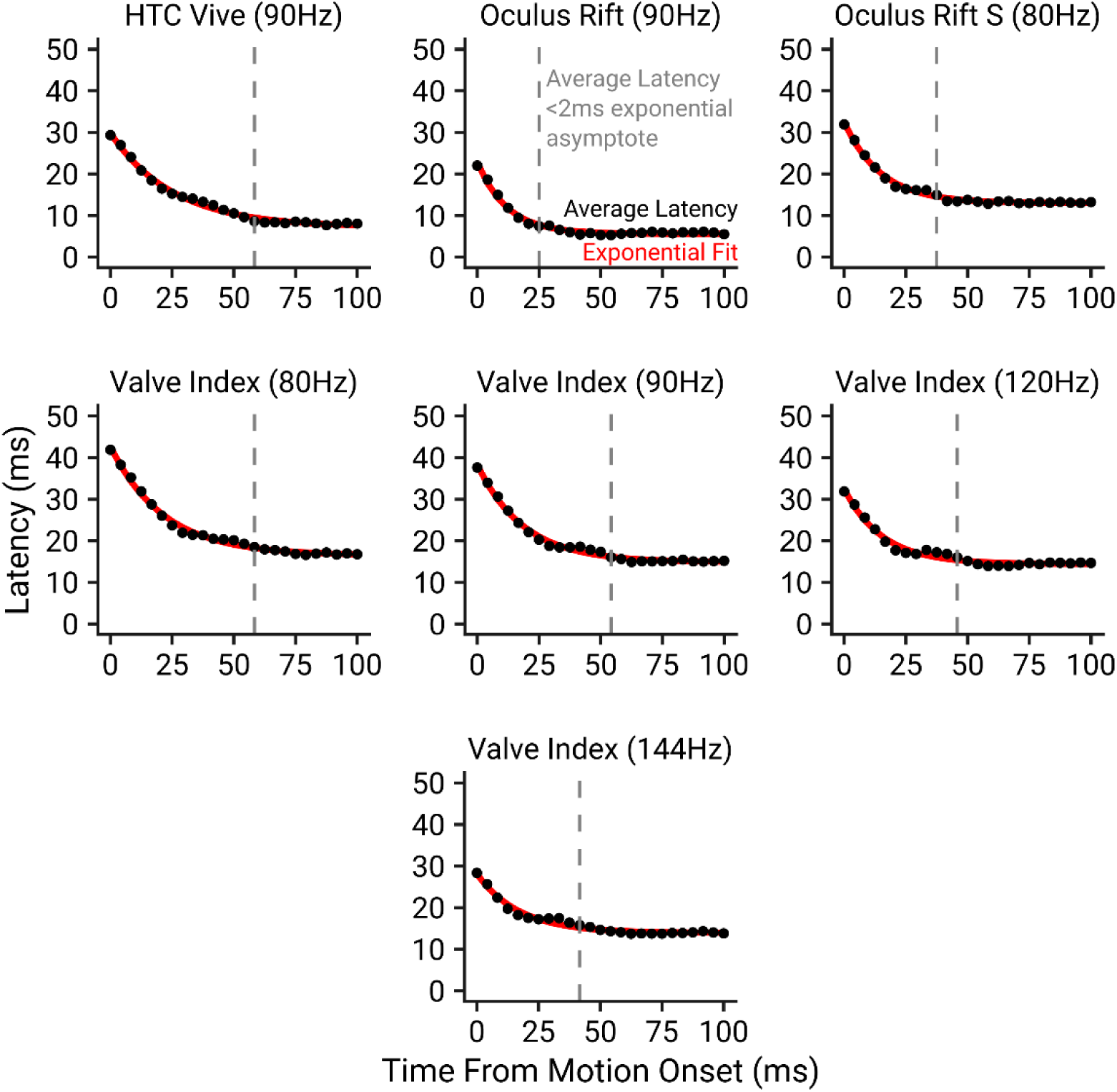
The minimum latency at which real controller positions could be displayed, as a function of the time from motion onset. The points show the average latency, the red curve shows the fit of an exponential function, and the dashed lines show the time at which the observed latency fell within 2ms of the exponential fit’s asymptotic latency.

While the latency can be interpreted literally as a temporal offset in positions, i.e. the virtual controller lags behind the real one, it may also be thought of as a positional offset, i.e. the virtual controller is always offset relative to the real one (Adelstein et al., 2003). To understand the implications of this, the real normalised controller position was taken from the virtual one at each time step, shown in Figure 5c. The low continuous latency of the Oculus Rift results in the average positional offset being negligible. While the Valve Index has a higher average positional offset, this is only 0.013 normalised units (∼2.4mm) when the latency has stabilised. These are, however, average values – the motion prediction will not work perfectly. Therefore, while the average temporal latency and positional offset are near-zero for the Oculus Rift, at any given time step these values are likely to be non-zero and will fluctuate.

## Discussion

While recent low-cost consumer VR HMDs have been rapidly adopted to study sensorimotor behaviour, these tools are not designed to be scientific instruments and as such, require verification to ensure the measurements they make are fit for purpose. Temporal latencies in the visual feedback of movement are known to degrade sensorimotor performance, but existing investigations test only HMD movements and use inconsistent software and hardware, making comparisons impossible and giving little idea of what latencies a typical research pipeline would show. We have developed a technique that allows researchers to measure the latencies throughout a movement and determine whether their experiment can be conducted in light of any technical limitations that might exist in VR hardware. We used this technique to measure the motion-to-photon latency of current popular HMD controllers (HTC Vive, Oculus Rift, Oculus Rift S, Valve Index). We found that (i) all of these HMDs show low levels of latency at the start of a movement (21-42ms); (ii) the latency stabilises near these lower values early in the movement (25-58ms from motion onset); and reduces further later in the movement once built-in motion prediction algorithms can be employed (2-13ms).

### Unity and SteamVR Timings

It is clear from the difference in latency between the sudden and continuous sections of the movement that motion prediction allows the latency to be functionally reduced. Using the Unity Profiler tool, which allows the user to see what functions were called on a frame and how long they took, the timeline of a frame being generated to being displayed on the HMD can be understood. A high-level overview of this timeline is shown in Figure 7a. Simulations are first run on the CPU, shown in Figure 7b, which includes running physics simulations and any user-defined logic in the Fixed Update, Update, and Late Update loops. Unity then waits until the end of the frame, just before rendering the frame, to allow SteamVR to update the tracking data (position and rotation) of any tracked objects. After this is done, the frame is offloaded to the GPU to be rendered to images, and then presented for display. When the rendered frame is presented, it is sent to the HMD and prepared to be shown. As low-persistence displays are used, the screens are illuminated for between 0.5-2ms (depending on system) at the end of the refresh cycle. These operations are aligned with the vertical-synchronisation of the HMDs, which maintains regular frame rates. The simulation, render, and presentation stages each take a frame to complete.

**Figure 7:**
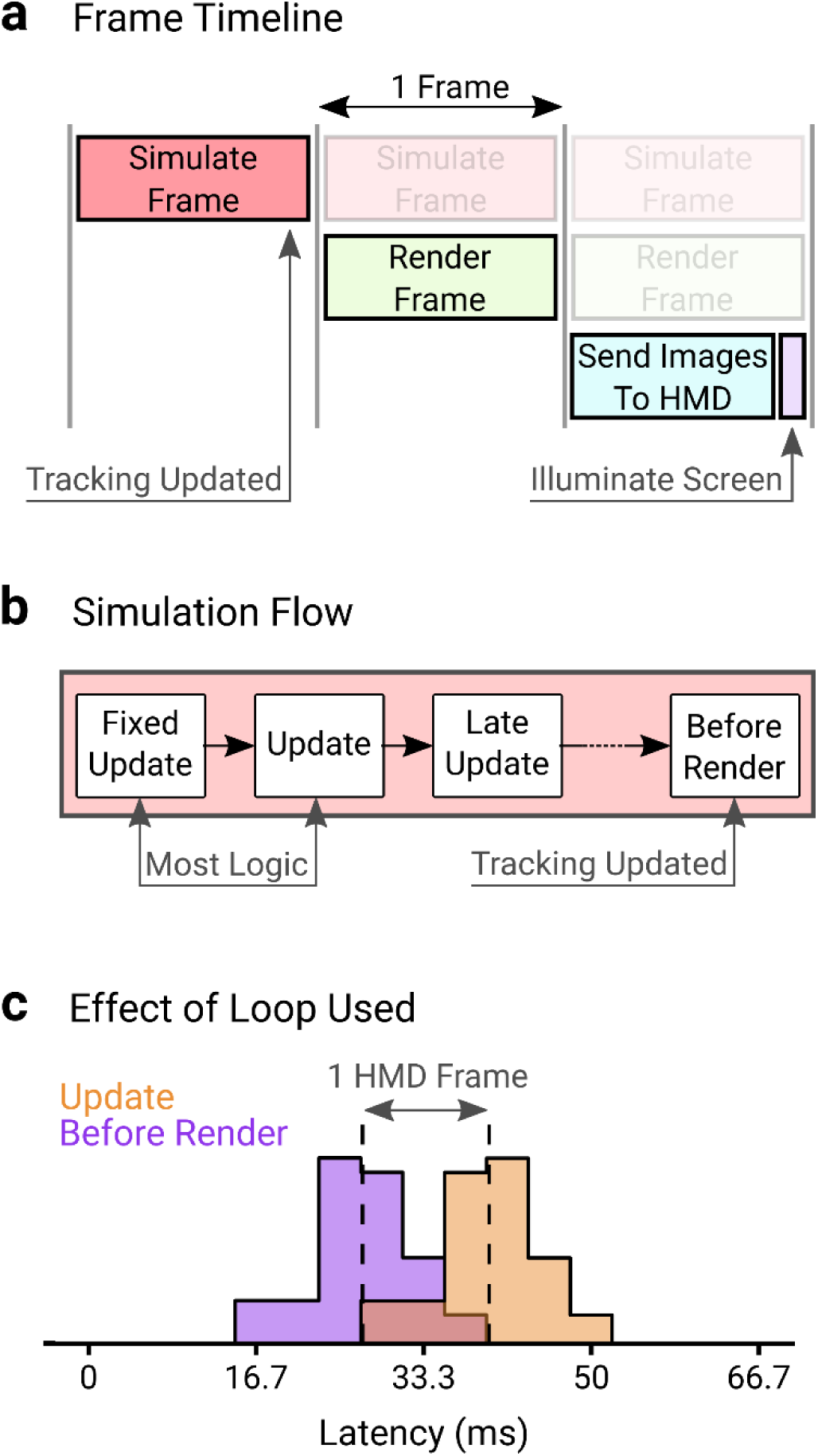
Unity + SteamVR frame processing operations **(a) Frame Timeline:** For a given frame to be shown to the user, it first must be simulated, rendered, sent to the HMD and then displayed. The tracking data (position and rotation) of tracked objects like the HMD and controllers are updated at the end of the simulation stage. **(b) Simulation Flow:** Inside the simulation stage, loops are called sequentially. First the Fixed Update loop operates, which handles physics operations, followed by the operation of the Update loop. Typically, logic for experiments (including game state changes, feedback, and interactions driven by controller movements) will be driven either by the Fixed Update or Update Loops. A Late Update loop operates, then there is a wait until the simulations are about to be rendered, where the Before Render loop runs, which is where the tracking data is updated. **(c) Effect of Loop Used:** The latency measurement technique was modified to detect virtual motion onset online using the same outlier detection algorithm, and the screens flashed to indicate movement had started. However, the screens were controlled in different loops, one in the Update Loop and one in the Before Render loop. As the tracking data are updated in the Before Render loop this reports motion earlier, whereas the Update loop captures motion onset one frame later, leading to a 1 HMD frame latency. The implication is using the position or rotation of the controller or HMD to drive logic will incur a 1 frame latency. The data shows one video for the Oculus Rift S.

**Figure 8:**
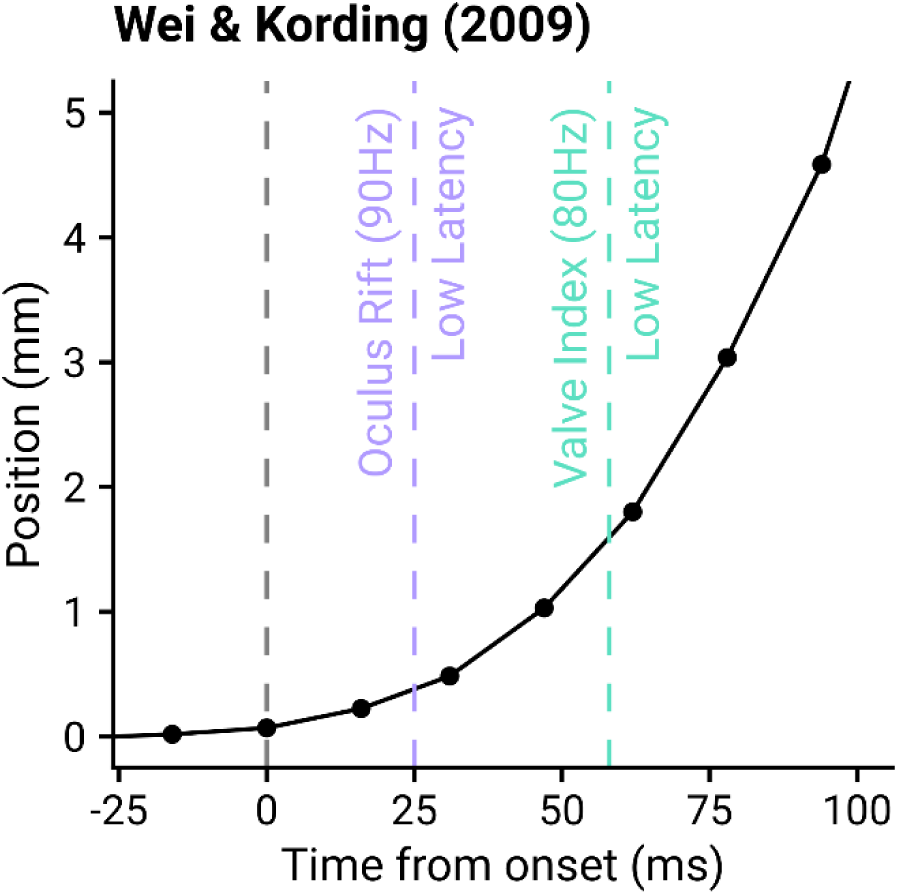
Data from Wei & Kording (2009), with lines indicating when the Oculus Rift (90Hz) and Valve Index (80Hz) would reach low levels of latency. The points and line show the average hand position in the x-direction across participants and trials, resampled to a fixed frame rate and aligned at motion onset, as identified according to Teasdale (1993).

Based on this high-level overview we should therefore expect the motion prediction to be predicting roughly two frames ahead, as the tracking data is updated at the end of the first frame of a three-frame process (Figure 7a). This is ratified by the SteamVR Unity plugin source code, making explicit that the predictions made are two frames ahead (SteamVR_Action_Pose.cs). That the latency is mostly ameliorated by motion prediction for the HTC Vive, Oculus Rift and Oculus Rift S suggests that the majority of the latency therefore comes during the rendering and presentation stages for these systems. Even after motion prediction, the Valve Index however has a significant amount of latency remaining which is consistent across frame rates. We surmise that the Valve Index has additional latency prior to the positions being sampled by SteamVR, or after display information is sent to the HMD by the graphics card. This is because there were no changes to computer or software, no issues in the SteamVR frame timing overlay, the Unity Profiler showed no difference in frame timelines during development across systems, and the Index’s remaining latency did not depend on the frame rate.

Because the equipment tracking data are updated in the Before Render loop, any physics operations or logic involving the position or rotation of the controller in the Fixed Update or Update loops will be working with the tracking data of the previous frame. While logic could be run in the Before Render loop, it is not advised by Unity as any additional work there will increase the visual latency. To demonstrate the effect of this, the Unity program was modified to turn the screen on for one frame when motion onset was detected (by implementing an online version of the motion onset detection used), and the left and right screen colours were updated in the Update and Before Render loops independently. Figure 7c shows that when the Update loop uses the position to drive logic it comes with an additional headset frame of latency. In an experimental context, this means that logic-based feedback (such as a target exploding when hit, text being displayed, or a sound playing) will be delayed by one frame.

In the past, the complexity of the scene being simulated and rendered may have also contributed to the observed latency. If completion of a step in Figure 7a immediately initiated the next step, then the time taken to perform these operations would add to the latency. However, this can lead to a visual artifact where parts of multiple frames are shown simultaneously on the display (Feldstein & Ellis, 2020). Modern VR systems, and often computers more generally, utilise vertical-synchronisation (VSync) to alleviate this problem, which presents a single rendered image to the display at fixed intervals. These VSync intervals are then used to synchronise each element of the pipeline, with the simulation and rendering steps each having a set amount of time to complete in. For simple scenes, most of the time per frame is actually spent waiting for the VSync interval, as shown in Figure S4 in Supplementary Materials. As long as the scene is not overly complex, such that the simulation or rendering steps overrun their allowed time, then the latency should not depend on the complexity of the scene being shown.

### Implications for sensorimotor experiments

For any movement with changes in direction, speed, or acceleration, the performance of the motion prediction algorithms, and hence the observed latency, will be dictated by how quickly changes in movement properties are being performed. If we consider the best method proposed in early research for the Oculus Rift (LaValle et al., 2014), the acceleration is assumed to be constant over the prediction interval (these principles will likely hold even if more sophisticated methods are used). Therefore, when the real controller motion is accelerating, the reported position will be underestimated, and when the real controller motion is decelerating, the reported position will be overestimated. However, the rate at which these changes occur in the experiment will dictate how successfully the movement will be represented (i.e. motion prediction will be less successful when changes occur very quickly). For an example of a typical sensorimotor reaching task, the data from Wei & Körding (2009) was collapsed across trials and participants to produce an average hand-position over time, aligned at motion onset as identified by the Teasdale method (1993). The points at which the latency became asymptotically low was marked for the best (Oculus Rift) and worst (Valve Index at 80Hz) performing systems, which indicated that across all tested systems participants would have moved by less than 2mm when latency reached low levels. In typical point-to-point movement tasks then, it is likely that the majority of the movement would be presented with low latency by a modern VR system.

Manual tracking tasks present useful case studies for considering the consequences of predictable and unpredictable movements on behaviour in VR. If the pattern being tracked is predictable, motion prediction should be able to ameliorate the latency through most of the movement. For example, in Smith (1972) participants were tasked with tracking a cursor moving along either a circular (20cm diameter) or octagonal (19cm square with short flattened corners) pattern at low speeds (120° per second), which should allow cursor motion to be predicted reasonably well. The study found that introduced delays degraded participant’s ability to stay on the target in a log-linear fashion, with even a 17ms delay degrading performance slightly relative to no delay. Therefore, it is likely that a small degradation in performance will be present when using off the shelf VR to study manual tracking relative to equipment with no delays (though this may be negligibly small). However, other manual tracking tasks employ unpredictable signals, such as the complex signals generated through combining different sine wave frequencies (e.g. Foulkes & Miall, 2000), requiring repeated changes of direction. If these changes in direction are performed quickly, the motion prediction algorithms would likely recognise the change in direction late and overshoot the stationary position slightly, effectively adding noise and increasing latency during the direction change. The extent of this overshooting behaviour will depend on the speed of direction changes, and future work should systematically assess the performance of systems across a range of movement frequencies. As this latency will be intermittent, and the range of additional latencies tested on these tasks is larger than the base latency for these systems (Foulkes & Miall, 2000; Miall & Jackson, 2006), it is unclear what effect this sort of overshooting behaviour might have on tracking behaviour. It is therefore important that researchers establish (and report) the nature of the impact of these delays on their experimental task.

Performance on another widely used paradigm in sensorimotor research, visuomotor adaptation, is also known to be degraded when delays are introduced. Kitazawa, Kohno & Uka (1995) had participants reach from a button to a target on a screen in front of them, with vision during the motion precluded and only displayed when the target was reached (terminal feedback). Prism goggles were used to displace vision to either the left or right, inducing motor adaptation to the displacement. Both the rate at which participants corrected the displacement while wearing goggles, and the initial error once the goggles were removed (indicating the amount of adaptation that occurred) were significantly degraded when the terminal feedback was delayed by 50ms or more, but not significantly when the feedback was delayed by 10ms or 20ms. However, it should be noted that these experiments were designed to assess the effect of delays over a large range (up to 10,000ms) rather than focusing on the small delays we saw in our study and were therefore likely underpowered to detect differences at 10 and 20ms if they did exist. Given that point-to-point reaches tend to be fairly smooth, where motion prediction should perform well, it is likely that both online and terminal feedback in visuomotor adaptation studies would be delivered with little latency. This implies there should be little degradation of motor adaptation because of the delays of the VR equipment, though it is possible a more highly powered study would find learning degradation at small delays.

Understanding the effects of small delays on adaptation would require further study. While other studies have investigated the effect of delays on adaptation, they often impose delays that are much larger than the base latencies of the VR systems considered (Brudner et al., 2016; Held et al., 1966; Schween & Hegele, 2017; Tanaka et al., 2011). Moreover, some setups have substantial base delays of ∼60ms (Honda et al., 2012), while others have utilised motion prediction algorithms (similar to those used in the VR systems studied here) to overcome their base latencies (Brudner et al., 2016; McKenna et al., 2017). This indicates that it is critical to know the latency inherent to the system being used, as the amount of adaptation observed will differ depending on whether a low or high latency setup was used.

### Comparison with other methods

The technique developed for this study was designed to overcome shortcomings of extant latency measurement methods. Many of the previously employed methods only give an idea of the latency at set points in a movement trajectory, usually either the start (Bryson & Fisher, 1990; Feldstein & Ellis, 2020; Raaen & Kjellmo, 2015; Seo et al., 2017; Yang et al., 2017), middle (Liang et al., 1991; Mine, 1993) or at some threshold point (Friston & Steed, 2014; He et al., 2000; Papadakis et al., 2011). We demonstrated that measures at a single point in the movement were not sufficient to capture the latency properties of current off the shelf VR systems because of motion prediction. Further, previous work that does compare the full real and virtual movement trajectory employ techniques such as cross-correlation to produce a single measure of latency (Adelstein et al., 1996; Becher et al., 2018; Di Luca, 2010; Gilson & Glennerster, 2012; Scarfe & Glennerster, 2019; Steed, 2008; Zhao et al., 2017). Again, this will not capture the latency properties of a system that uses motion prediction well, as these measures will likely be weighted heavily towards the ‘continuous’ latency. Instead, we have developed a technique that can provide a measure of the latency through the whole movement to demonstrate the effect of motion prediction.

Furthermore, we wanted our technique to be user-friendly and accessible to any researcher wanting to measure their own system. Many previously developed techniques that assess full motion profiles use electrical hardware that are not easily deployed in many sensorimotor laboratories – such as rotary encoders, motors, photodiodes, amplifiers and oscilloscopes (Adelstein et al., 1996; Becher et al., 2018; Di Luca, 2010; Zhao et al., 2017). The advantage of our technique is that all of the mechanical hardware required can be bought or fabricated relatively cheaply, and the only additional electrical hardware used is a smart phone capable of recording at 240fps or above. While existing techniques use a high-speed camera to measure latency (Gilson & Glennerster, 2012; Steed, 2008), it requires the virtual representation be visible to the camera. This would be difficult for HMDs, as any representation will likely be too small to be measured accurately, with additional warping due to the lenses. Our method does not attempt to represent the virtual motion parametrically, but rather uses the colour of the HMD’s screen to co-register the real movements, captured by the camera, and the virtual movement, written to file with the corresponding screen colour.

The presented technique could be expanded upon by assessing the response of the systems to movements at different frequencies. While the current technique allows the effect of motion prediction to be characterised during simple movements, it is unclear how fast participants would have to move for the motion prediction to perform badly, for example when making changes of direction in a manual tracking task. This could be assessed by linearly actuating the controller at known frequencies, and assessing the gain and latency response of the system (e.g. Adelstein et al., 1996). Knowing the system’s response across a frequency range, and comparing that top the frequencies with which people can move their arms, would fully provide further insight into the types of movements that can be reliably measured using current VR systems.

## Conclusion

We have created a novel technique to measure the motion-to-photon latency of controller movements. We tested the technique using four popular VR HMDs – the HTC Vive, Oculus Rift, Oculus Rift S, and Valve Index. Measurements of latency were made at the start of the movement, where all latencies were between 21-42ms on average, and at the middle of the movement, where all latencies were 2-13ms on average. Measurements of latency through the whole movement showed the motion prediction algorithms reduce this latency within the first 25-58ms of the movement. There are likely to be a number of experimental tasks and research questions where these latencies (and the variability thereof) are not problematic. Conversely, there will be tasks that are adversely affected by these temporal delays and the fact that the latencies vary throughout the movement. It is therefore incumbent on researchers to determine the latencies associated with the equipment they are using, report these latencies, and ensure that conclusions drawn from their experimental results are not undermined by this potentially confounding variable. The technique we report within this manuscript (and the associated software that we have made freely available) will enable researchers to make these measurements. In turn, this should enable the potential of VR systems in research to be realised.

## Availability

All scripts required to perform and analyse the latency measurements will be made available upon publication at the Open Science Framework.

## Acknowledgements

Authors F.M and M.M-W were supported by Fellowships from the Alan Turing Institute and a Research Grant from The Engineering and Physical Sciences Research Council (EPSRC) (EP/R031193/1).

