## Supplementary Materials for "Measuring motion-to-photon latency for sensorimotor experiments with virtual reality systems"

#### Ambiguous Screen Colour Handling

Due to the camera's rolling shutter, some camera frames captured the HMD screen just turning on or off. When this happened, it meant there were few pixels inside the ROI on which to base the colour classification for the screen, which we found to be less accurate. However, the consistency of the direction of the rolling shutter meant knowing whether those few pixels inside the ROI were at the top or bottom could be leveraged to correct the screen colour post-hoc during analysis. When the small number of illuminated pixels were featured at the top of the ROI, the frame prior must have been illuminated, illustrated in Figure S1. Because both frames were the same colour, this must have represented a single illuminated frame, otherwise the colour on the screen would have changed. The opposite is also true – if the small number of illuminated pixels were featured at the bottom of the ROI, the following frame must feature the ROI fully illuminated. These frames where the ROI is only partially illuminated can then be corrected accurately in post-hoc analysis.

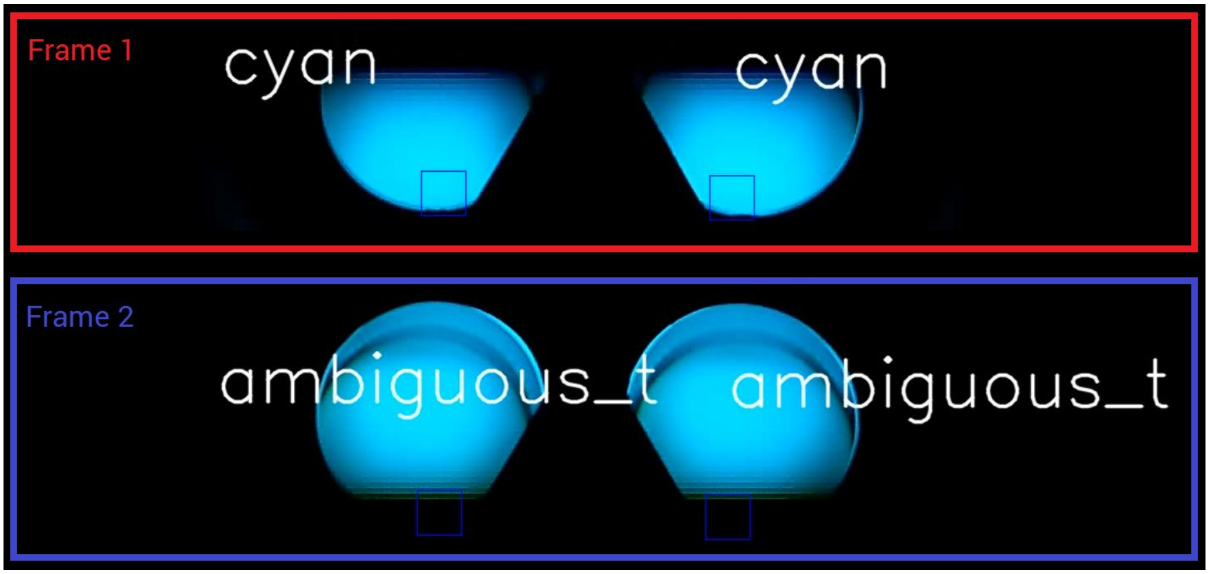


***Figure S1:*** An example of an 'ambiguous' frame. On Frame 1 the camera captures the display in the process of being illuminated. The rolling shutter of the camera means the top was captured while the screen was still off, so the top half of the screen is blank. Then in Frame 2 the camera captures the display in the process of being turned off. The rolling shutter means this time the top of the display is still on but the bottom has turned off. The second frame was marked as being ambiguous due to low pixel count in the ROI. We can see that because both frames feature the same colour it must be from the same HMD frame (otherwise the screen colour would have changed). Hence the surrounding frames can be used to accurately fill in 'ambiguous' frames.

### Latency Measurement Variability

Some of the variability in latency measurements presented in Figure 4b arise due to the definition of latency used. For both measures presented in 4b, the difference in time between two discrete events is measured – either the difference in time between the real and virtual controller starting to move or crossing the mid-point of the movement. Different latency measurements can arise due to the stochasticity between the time at which the event occurs, and when the camera frame is captured and the HMD displays a new frame, as illustrated in Figure S2.

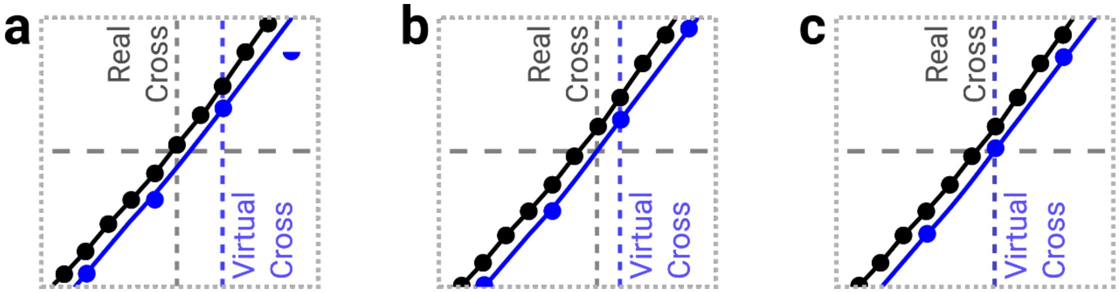


***Figure S2:*** Three possible ways that the same real and virtual controller positions could fall such that the cross-correlation of the lines is the same, but the measured latency is different, showing (a) a two-frame latency, (b) a one-frame latency, and (c) a zero-frame latency.

### Overshooting Behaviour

A logical consequence of the motion prediction system used is that when the real controller suddenly comes to rest, the virtual controller will overshoot before correcting for this sudden change in speed. The same procedure used to detect motion onset was performed in the reverse direction to detect motion offset, and the controller trajectory was then time-normalised about the offset. This can be seen for the Oculus Rift in Figure S3. On average the Oculus Rift controller starts to correct for the overshoot after around 5 camera frames (20.8ms, similar to the sudden start latency measured for the system). While the sudden start of the movement was applied consistently, the end of the movement was relatively less consistent between movements, and also between systems, meaning that comparisons of overshooting behaviour between systems is not informative. Future work could assess this overshooting in more controller movements, for example by using an electric motor to translate the assembly in a reliable manner.

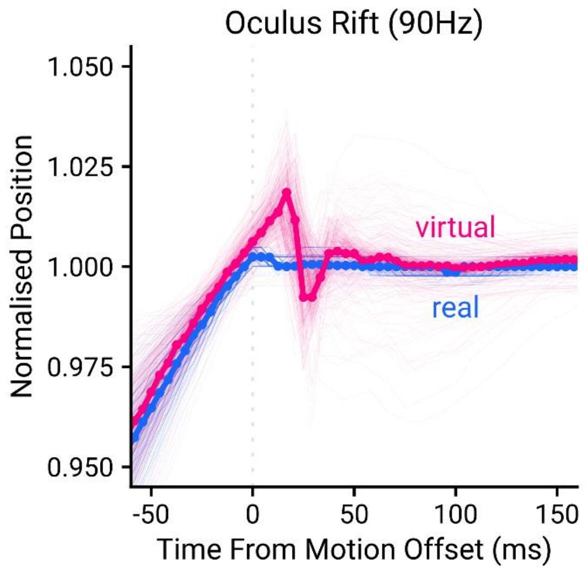


***Figure S3:*** Real and virtual controller trajectories across time, aligned at motion offset. The virtual controller shows overshooting upon a sudden stop, a logical consequence of the use of motion prediction.

### Frame Timings, Scene Complexity, and Latency

Figure S4 shows an example of the pipeline by which a frame is simulated and rendered in the Unity Profiler. The “Main Thread” simulates the scene, performing operations like physics updates, changes to the scene appearance, etc. The render thread then renders the scene. Both steps are aligned with the VSync interval of the headset. This shows an example from the scene used to perform the latency measurements.  Notably, most of the simulation and render steps are spent waiting for the VSync interval – each thread is given a set amount of time to perform operations within to maintain synchronisation of the pipeline. As long as the scene is complex enough to cause either thread to overrun their allowed time, the complexity of the scene being shown should have no effect on the observed latency.

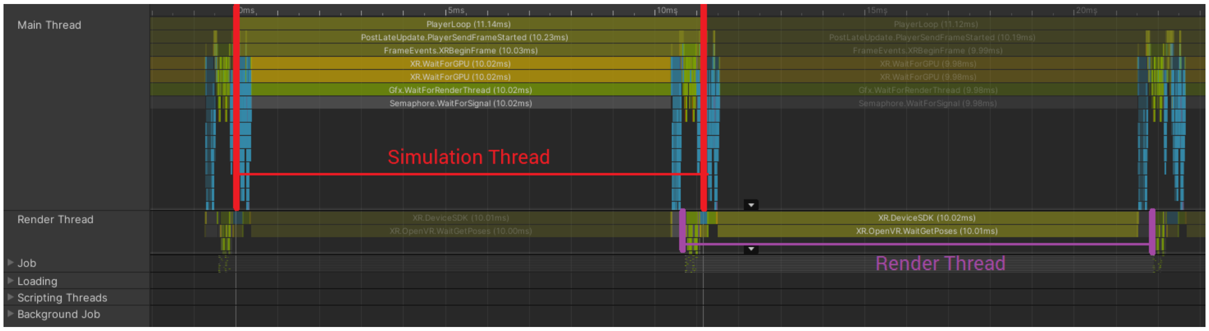


***Figure S4***: Frame timings from the Unity Profiler. The Simulation and Render thread are aligned with the VSync intervals, giving each thread a set amount of time to perform all of the necessary steps. In this example, most of the frame is spent waiting to perform the rendering commands, so more complex scenes could be rendered with no hit to the latency.
